## Supplementary material for "The effect of temperature on the boundary conditions of West Nile virus circulation in Europe"

### Supplementary Text 1

We obtained human reported cases for the European Union from the European Surveillance System (TESSY). In the Nomenclature of Territorial Units for Statistics (NUTS) 2 regions of Utrecht (NL31), Berlin (DE30), Brandenburg (DE40), Andalucía (ES61), Veneto (ITH3), Emilia-Romagna (ITH5), Kentriki Makedonia (EL52), 1443 (58%) out of the total of 2487 human West Nile fever cases were reported to TESSY between 2010 and 2021 (Fig S2.1 and S2.2).

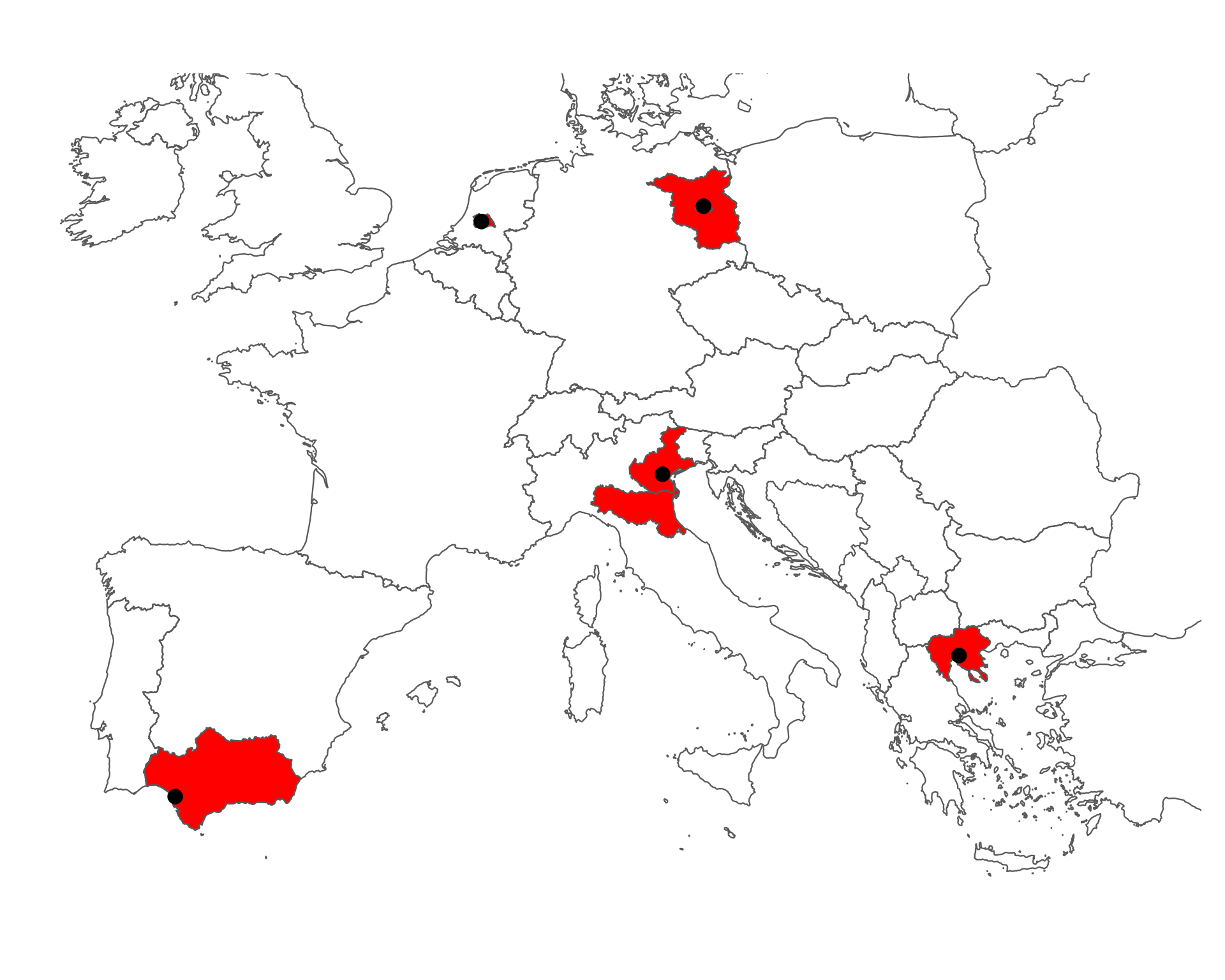

**Fig S1.1:** Location of the selected locations (points) and highlighted the surrounding Nomenclature of Territorial Units for Statistics (NUTS) 2 regions of Utrecht (NL31), Berlin (DE30), Brandenburg (DE40), Andalucía (ES61), Veneto (ITH3), Emilia-Romagna (ITH5), Kentriki Makedonia (EL52).

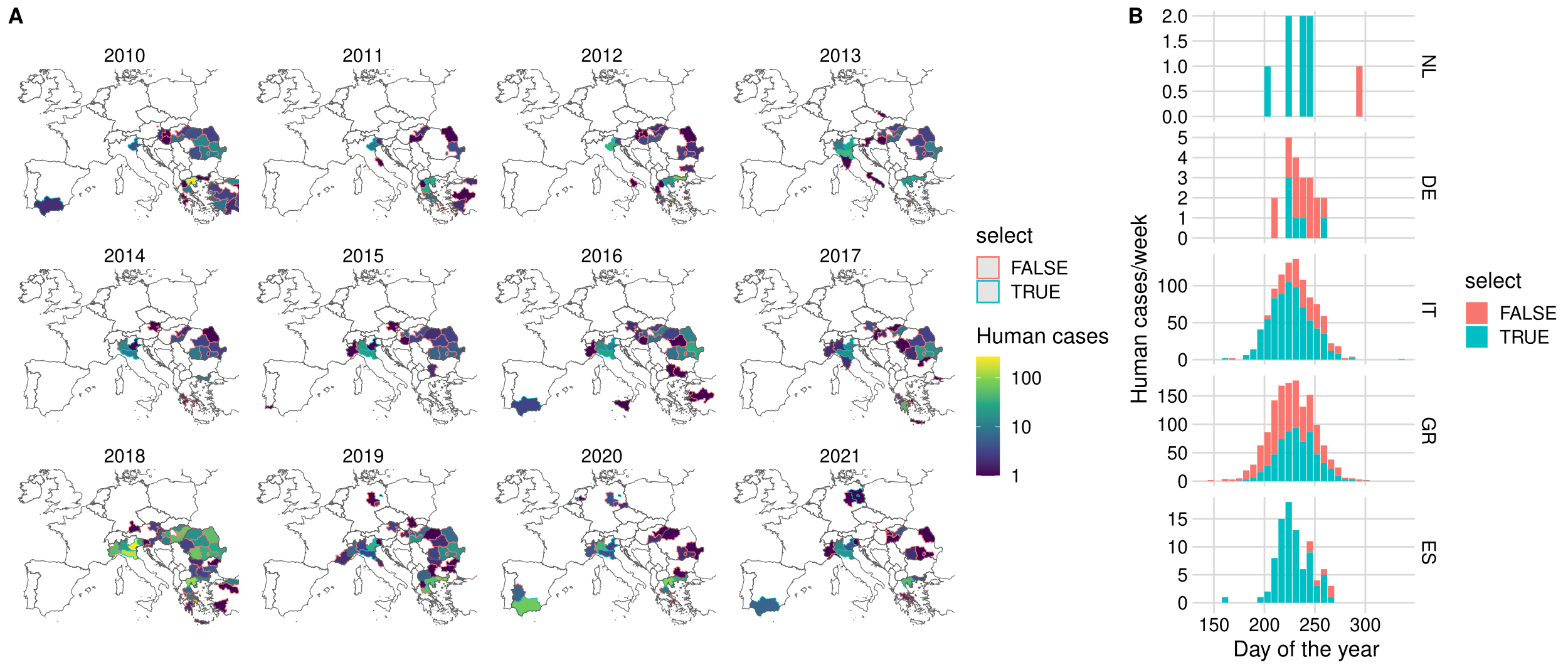

**Fig S1.2:** A. B. Cumulative number of weekly human West Nile cases for the period 2010-2021 for the five countries. Colour indicates the proportion that was in the NUTS2 regions for which we assessed the boundary conditions. The Netherlands [NL], Germany [DE], Italy [IT] and Greece [GR], Spain [ES].

### Supplementary Text 2

Equation S1 (quadratic) and S2 (Briére) are functions of temperature (*T*). These depend on a lower and upper thermal limit (*T_min,_,T_max_*) and a coefficient (*q*).

Quadratic: $f(T)=-q(T-T_{min})(T-T_{max})$	(S1)

Briére: $f(T)=q\cdot T(T-T_{min})\sqrt{(T_{max}-T)}$	(S2)

These functions were fit to laboratory observations where under controlled temperature regimes disease and lifecycle parameters were estimated (see Fig 2).

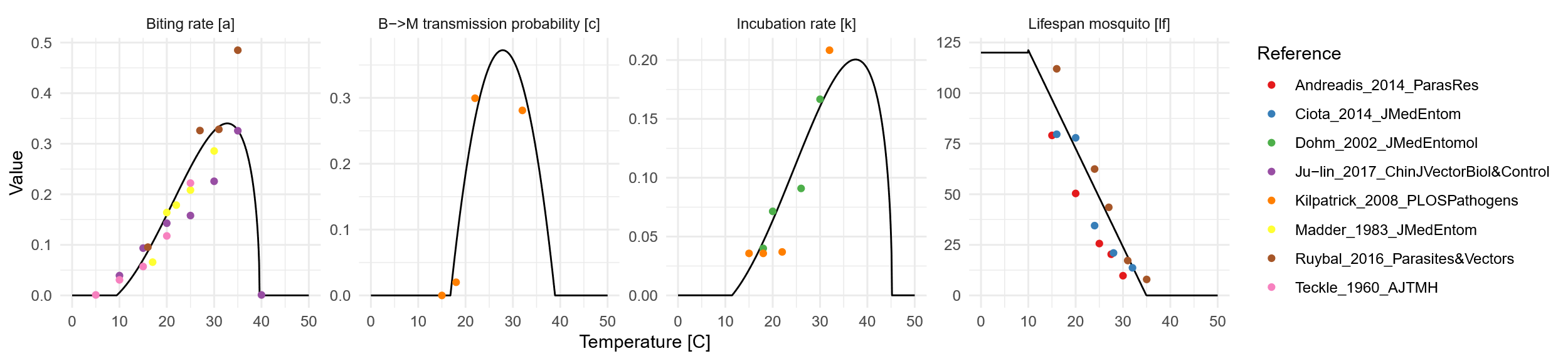

**Fig S2.1:** Temperature-response curves on mosquito biting rate (*a*, days^-1^), proportion of mosquitoes that become infectious after taking a blood meal on an infected bird (c), rate at which mosquitoes become infectious (*k* days^-1^, 1*/k* = incubation time in days), and mosquito lifespan in days (*lf*, *µ_A_* = 1/*lf*).

### Supplementary Text 3

To assess the West Nile virus (WNV) prevalence in mosquitoes in endemic regions in Spain, Italy and Greece, we collected data from literature. We included studies where the presence of WNV was assessed in *Cx. Pipiens* mosquitoes, and that were conducted from 2010 until 2022, the year of the literature search. We identified 16 studies that qualified these inclusion criteria [1-16]. In these studies mosquitoes were trapped and pooled in groups of 50-200 mosquitoes, and occasionally in smaller pools. We extracted the number of positive pools, the number of pools tested, the pool size and the total number of mosquitoes tested. We calculate both the minimum infection rate (MIR), as the ratio of the number of positive pools (Y) to the total number of mosquitoes tested (t), and the MLE (Equation S1), with the number of positive pools (Y), the number of pools (X), and the pool size (m) assuming a constant pool size [17]. Since often the exact pool sizes were not reported, we assumed a constant pool size calculated by the reported number of tested mosquitoes divided by the reported number of pools tested. When infection rates are higher, these methods are known to underestimate the infection rates. Furthermore, we calculated the average weekly trapping rate by dividing the total number of trapped mosquitoes by the number of trapping weeks.

$MLE=1-{(1-\frac{Y}{X})}^{1/m}$	Eq. S1

We found in these areas that on average 1.0/1000 (MIR) or 1.2/1000 (MLE) of the trapped *Cx Pipiens* mosquitoes were positive. The median weekly trapping rate was 955 mosquitoes, but varied greatly between studies since the number of traps used varied similarly.

**Table. S1. Summary of mosquitoes studies conducted in Italy, Greece and Spain between 2010-2022.**

| **Country** | **Year(s)** | **Trapping months** | **Mosquitoes tested** | **Pools test (positive)** | **MIR^a^** | **MLE^a^** | **Trapping**  **Rate (week^-1^)** | **Reference** |
| --- | --- | --- | --- | --- | --- | --- | --- | --- |
| Italy | 2018 | 5 | 269112 | 2331 (232) | 0.86 | 0.91 | 11961 | [52] |
| Italy | 2008-2014 | 8 | 2346224 | 106 (5) | 0 | 0 | 65173 | [53] |
| Italy | 2010-2011 | 7.5 | 1156 | 131 (0) | 0 | 0 | 34 | [53] |
| Italy | 2014 | 5 | 10757 | 914 (3) | 0.28 | 0.27 | 478 | [54] |
| Italy | 2014 | 5 | 78131 | 1514 (9) | 0.12 | 0.11 | 3472 | [55] |
| Italy | 2013 | 5 | 510773 | 7256 (178) | 0.35 | 0.35 | 22701 | [56] |
| Italy | 2010-2012 | 18 | 262905 | NR (23) | 0.09 | NR | 3246 | [57] |
| Italy | 2011 | 5 | 69025 | 2732 (5) | 0.07 | 0.07 | 3068 | [58] |
| Italy | 2009-2011 | 2 | 4799 | 172 (0) | 0 | 0 | 533 | [59] |
| Italy | 2008-2011 | 5 | 818474 | 5645 (32) | 0.04 | 0.04 | 36377 | [59] |
| Italy | 2011 | 5 | 69172 | 2732 (5) | 0.07 | 0.07 | 3074 | [60] |
| Greece | 2020 | 4 | 2809 | 70 (10) | 3.56 | 3.85 | 156 | [61] |
| Greece | 2018 | 5 | 17470 | 229 (10) | 0.57 | 0.59 | 776 | [62] |
| Greece | 2010 | 2 | 6597 | 110 (2) | 0.3 | 0.31 | 733 | [63] |
| Greece | 2011 | 6 | 61812 | 824 (60) | 0.97 | 1.01 | 2289 | [63] |
| Greece | 2012 | 5 | 87643 | 663 (98) | 1.12 | 1.21 | 3895 | [63] |
| Greece | 2013 | 5 | 3906 | 130 (44) | 11.26 | 13.68 | 174 | [63] |
| Greece | 2013-2013 | 6 | 25780 | 295 (9) | 0.35 | 0.36 | 955 | [64] |
| Greece | 2010-2011 | 8 | 16116 | 296 (6) | 0.37 | 0.38 | 448 | [65] |
| Greece | 2010 | 1 | 3232 | 65 (2) | 0.62 | 0.62 | 718 | [66] |
| Spain | 2020 | 6 | 1563 | 152 (1) | 0.64 | 0.66 | 58 | [67] |

^a^ minimum infection rate (MIR) and maximum likelihood estimation (MLE) of the infection rate per 1000 mosquitoes. NR = not reported/unable to calculate due to missing data.

#### References S3 Text

### Fig S1. The proportion of introductions that result in a West Nile virus outbreak in The Netherlands, Germany, Italy, Greece and Spain, assuming a fully susceptible population in 2020.

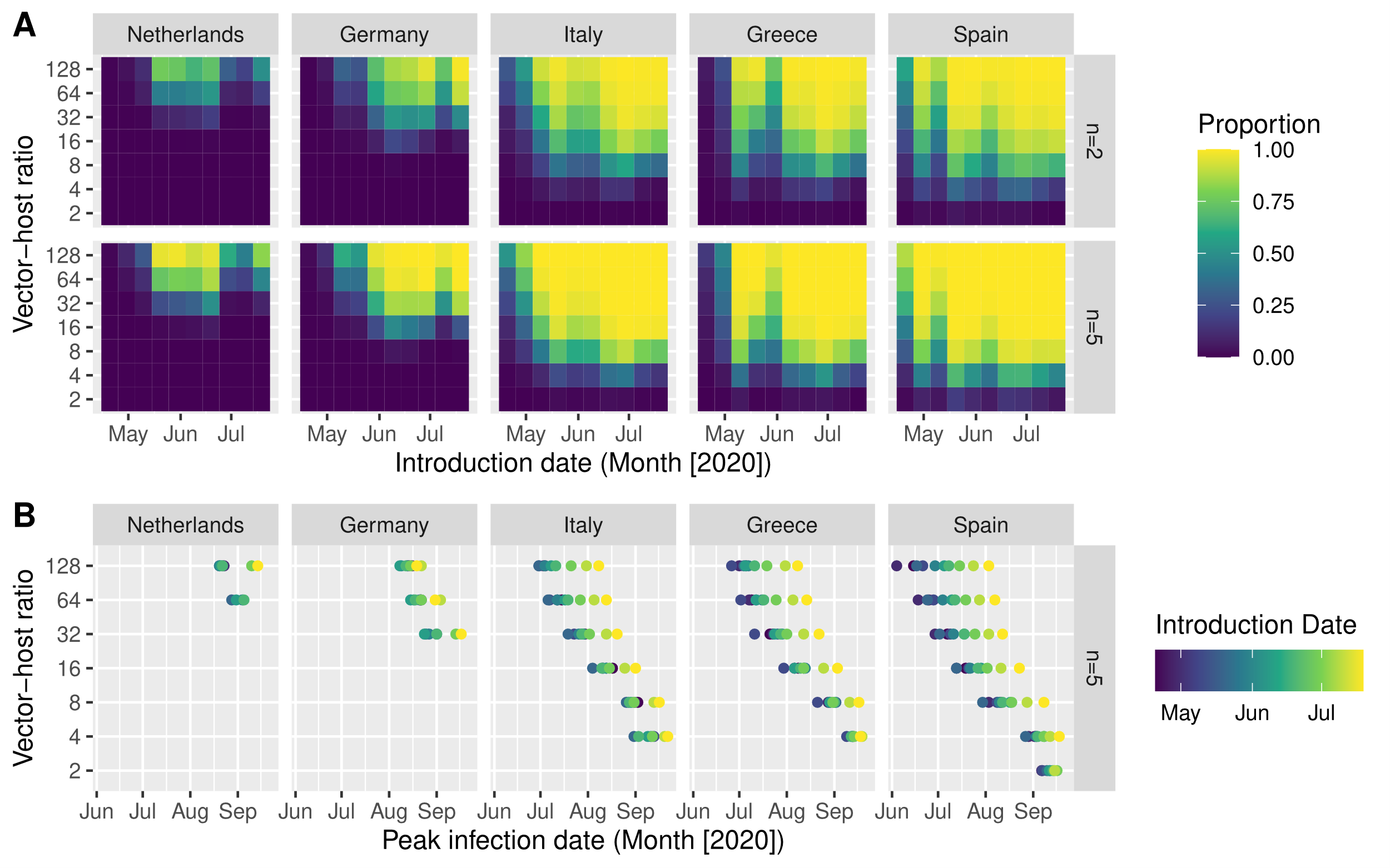

**Fig S1:** **A.** The proportion of introductions that result in a WNV outbreak by introduction date (x-axis) and vector-host ratio (y-axis); for five locations (facet columns) and with an introduction of 2 or 5 of infected birds (facet rows, n=2, n=5) **B.** Date on which the proportion of infected mosquitoes surpasses 1% (x-axis) after introductions at different vector-host ratios (y-axis) and introduction dates (colour) for five locations (facet columns). Locations: The Netherlands (Utrecht), Germany (Berlin), Italy (Po-valley) and Greece (Aksiou Delta), Spain (Doñana National Park).

### Fig S2. The proportion of introductions that result in a West Nile virus outbreak in Berlin between 2011-2022.

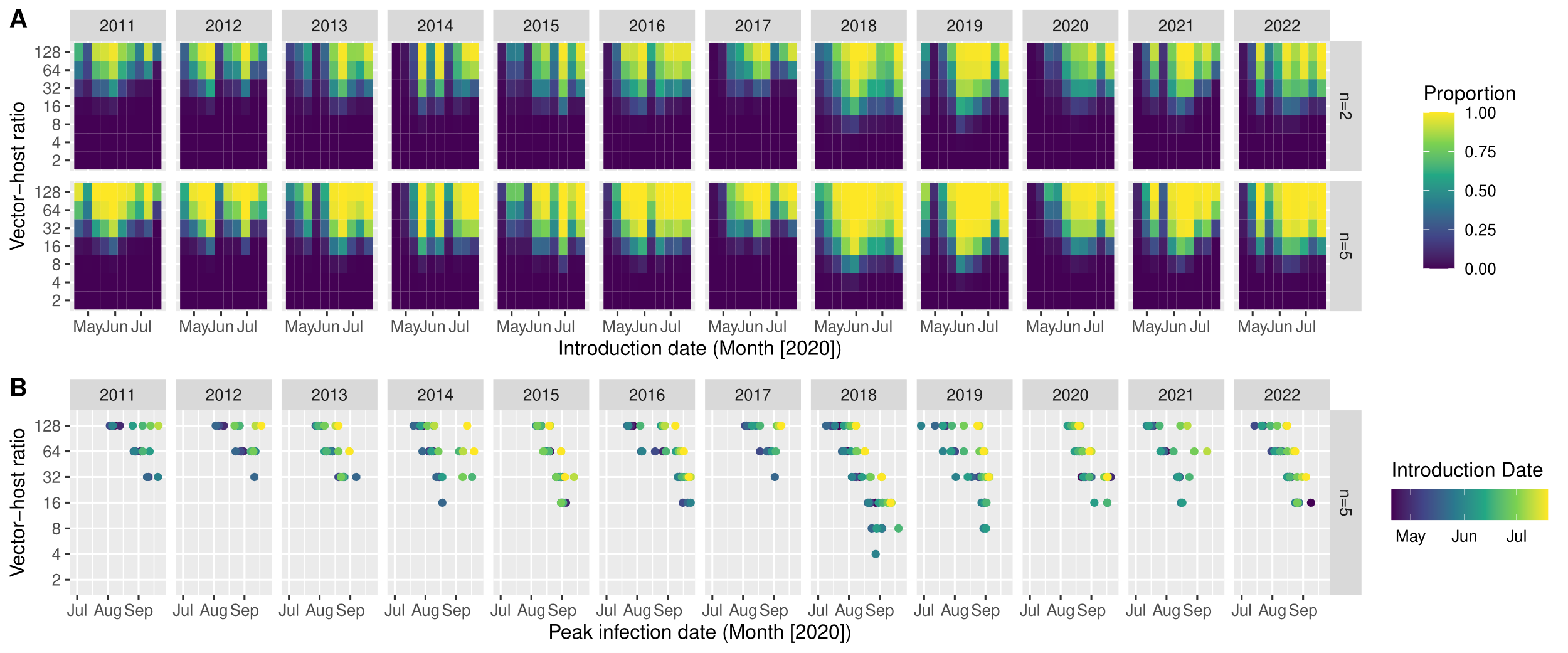

**Fig S2:** **A.** The proportion of introductions that result in a WNV outbreak by introduction date (x-axis) and vector-host ratio (y-axis); for five locations (facet columns) and with an introduction of 2 or 5 of infected birds (facet rows, n=2, n=5) in Germany (Berlin) **B.** Date on which the proportion of infected mosquitoes surpasses 1% (x-axis) after introductions at different vector-host ratios (y-axis) and introduction dates (colour) for Germany (Berlin) by year (facets).
